# Evaluation of gaseous ozone technology for decontaminating porcine reproductive and respiratory syndrome virus (PRRSV) and porcine epidemic diarrhea virus (PEDV) from non-porous surface in truck cabins

**DOI:** 10.64898/2026.07.23.740241

**Authors:** Thinh Tran-Pham-Tien, Jianqiang Zhang, Mafalda Pedro Mil-Homens, Ethan Aljets, Hao Tong, Baoqing Guo, Kate Dion, Peng Li, Mariáh Post, Elly Kirwa, Elisa De Conti, Daniel C. A. Moraes DC, Guilherme A. Cezar, Isadora Machado, Onyekachukwu Henry Osemeke, Gustavo Silva, Giovani Trevisan, Daniel Linhares, Derald J. Holtkamp

## Abstract

Porcine reproductive and respiratory syndrome virus (PRRSV) and porcine epidemic diarrhea virus (PEDV), both economically detrimental to the swine industry, can be spread through contaminated truck cabins, posing significant transport biosecurity risks. This study evaluated the effectiveness of gaseous ozone technology in decontaminating PRRSV and PEDV on non-porous surfaces in truck cabins. An incomplete factorial study comprising 26 treatment groups, including controls, and ozone treatments at three capacities (30, 38, and 68 g/h) was conducted over varied exposure times (0.5, 1, or 2 hours) in a climate-controlled truck cabin, where PRRSV or PEDV contaminated rubber coupons were subjected to treatments under continuous environmental monitoring, followed by virus elution and titration in cell cultures. Statistical analyses included regression models and ANOVA with Tukey’s comparisons, standard deviations for ozone machines, and Pearson correlations between ozone concentration and environmental factors. Ozone treatments showed variable results with less than two-log viral titer reduction. Ozone concentrations varied across identical setups (SD: 4.75 ppm for 30 g/h, 5.43 ppm for 38 g/h machines) during the first 30 minutes, while across the entire study period, ozone concentration showed weak negative correlation with virus titer (p > 0.05), moderate positive correlation with temperature (r = 0.5, p < 0.0001), and moderate negative correlation with humidity (r = -0.56, p < 0.0001). In conclusion, ozone treatments showed inconsistent and limited effectiveness in PRRSV and PEDV decontamination, influenced by temperature and humidity under the tested conditions.

## Introduction

Porcine reproductive and respiratory syndrome virus (PRRSV) and porcine epidemic diarrhea virus (PEDV) are contagious viral agents that have a significant economic impact on the swine industry (1,2). PRRSV alone is estimated to cause annual losses nearly doubling to $1.2 billion in the United States (3), while PED outbreaks in 2013-2014 resulted in the loss of approximately 7 million piglets (4). These viruses can be transmitted through various routes, including contaminated vehicles such as truck cabins, posing a biosecurity risk (5,6). Given the evidence of the potential for virus spread through contaminated vehicles (7,8), it is crucial to identify effective methods for preventing and mitigating the risk of virus transmission in the context of truck transportation in the swine industry.

Several decontamination processes for truck cabins are currently available, including the use of chemical disinfectants (9,10), heat treatment (11,12), and ultraviolet light (13,14). However, these methods have some drawbacks, such as the potential for harmful chemical residues (15), the time required for heat treatment (16), and the limited penetration of ultraviolet light (17). As a result, there is a need for alternative decontamination methods that can effectively inactivate viruses while minimizing these drawbacks.

Ozone technology is a promising alternative method for air and surface decontamination by inactivating viruses through oxidation (18–21). In recent years, particularly in the wake of the COVID-19 pandemic, there has been a growing interest in the development and use of ozone gas for air and surface decontamination (22–25). Ozone technology in its gaseous form can spread thoroughly throughout the entire space, with the ability to permeate, perfuse, and disinfect the “nooks and crannies” within enclosed spaces (26,27).

Ozone (O_3_) is a powerful oxidant that can inactivate viruses through two pathways: direct inactivation, involving the immediate oxidizing impact of ozone, and indirect inactivation, occurring through the reaction of liberated hydroxyl radicals arising from ozone decomposition, both of which damage the structural integrity of bacteria and viruses, targeting viral capsids and RNA, disrupting their reproductive cycle and ultimately leading to their elimination (18,28–30). Although the virucidal activity of ozone against various viruses has been studied (18,31), its specific effects on PRRSV and PEDV at different concentrations and exposure times on surfaces in truck cabins have not been investigated.

The objective of this study was to evaluate the efficacy of gaseous ozone technology for inactivating PRRSV and PEDV on non-porous surfaces found in truck cabins. By addressing this objective, the study seeks to provide insights into the adoption of new practical decontamination strategy to reduce the risk of virus transmission via trucks in the swine industry.

## Materials and methods

### Study design

This study employed an incomplete factorial design to evaluate the impact of alternative treatment types and exposure times on the inactivation of PRRSV and PEDV on rubber coupons. The factors included three exposure times (30, 60, or 120 mins), three treatment types (Ozone 30 g/h, Ozone 38 g/h, and Ozone 68 g/h), and untreated positive controls for each virus (Table 1). The three treatment types were studied using two commercially available ozone machines (TTDMK Ozone Generator, Flushing, New York) with ozone generation capacity of 30 and 38 g/h. A single ozone machine was used for the 30 and 38 g/h ozone treatments, while the 68 g/h ozone treatments comprised a combination of two ozone machines (30 and 38 g/h).

**Table 1.**
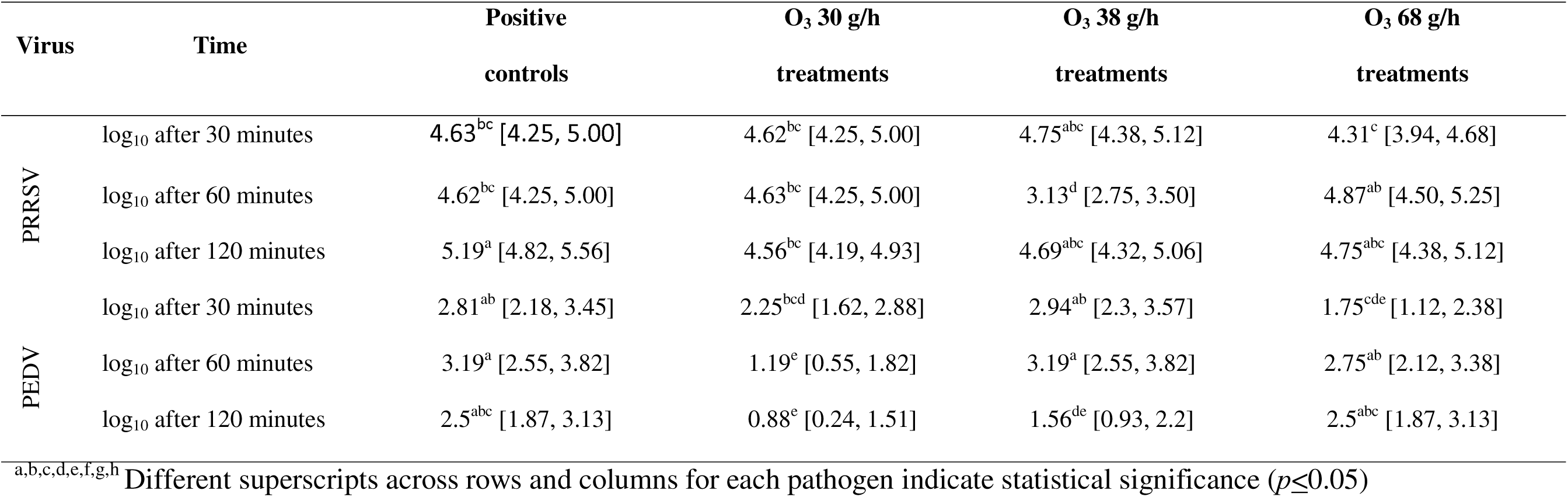
Least-squares means viral titers expressed as the log_10_TCID_50_/ml with 95% confident interval for each treatment group.

In total, with two viruses, there were 26 treatment groups with 4 replicates each, including 24 treatment groups described in Table 1 and two negative control treatment groups (one for each virus). Each negative group consisted of uncontaminated coupons subjected to the 120-minute exposure time without exposure to any virus or decontamination equipment. For positive controls, coupons contaminated with each virus were evaluated at each exposure time, resulting in 6 distinct positive control treatment groups. The ozone treatment groups comprised 18 groups, representing three different ozone generator capacity levels (30 g/h, 38 g/h, or 68 g/h) across three exposure times (30, 60, or 120 minutes) for each of the two viruses.

This comprehensive design allowed for rigorous assessment of viral inactivation efficacy under the various experimental conditions. All experimental protocols were carried out following approval from Iowa State University’s Office for Research Ethics (IBC Log #19–085). Procedures to contaminate the coupons were performed in a BSL-2 facility equipped with biosafety cabinets to prevent aerosol cross-contamination between treatment groups.

### Virus isolates

Two viral isolates were used in this study. The PRRSV 1-4-4 L1C.5 USA/MN/01775GA/2021 isolate (Rawal et al., 2023) was propagated in MARC-145 cells as outlined by Neat et al. (2021), with a resulting viral titer of 6.70 log_10_TCID_50_/ml. PEDV USA/NC49469/2013 P8 (Chen et al., 2016) was propagated in Vero cells following the protocol described by Chen et al. (2014), resulting in a final titer of 6.30 log_10_TCID_50_/ml.

### Coupon construction and contamination

Rubber coupons were constructed using material similar to that used to manufacture floor mats in vehicles. The coupons were cut to produce a circular coupon that was approximately 8.9 centimeters in diameter. Four coupons were randomly allocated to each treatment group and placed in a single sealed bag for autoclaving before inoculation. The contamination process was conducted inside separate biosafety cabinets for negative controls, positive controls, and ozone treatments to minimize cross-contamination risk. PRRSV and PEDV inoculations were performed on separate days to further reduce the risk of viral cross-contamination. The biosafety cabinet was decontaminated between each treatment group by wiping the surfaces with a 70% isopropyl alcohol solution followed by a disinfecting wipe (Clorox, Clorox Company, Oakland, California).

Each coupon was placed inside a separate petri dish measuring 100 mm in diameter and 15 mm in height (Fisherbrand Petri Dishes with Rimless Lid, Thermo Fisher Scientific, Waltham, Massachusetts). Negative control coupons were sham-contaminated with 2 ml of Minimum Essential Medium (MEM; SIGMA, United States). All other treatment groups were contaminated with 2 ml of PRRSV or PEDV stock solution. The virus stock solution or MEM was pipetted onto the upper surface of each coupon, with care not to allow any liquid to run off the edges. A new pipette tip was used to scrape and disperse the inoculum evenly over the top surface of each coupon. The inoculated coupons were left to air-dry inside the biosafety cabinet for 2 hours. At the end of the drying time, the lid was placed on each petri dish and placed in aluminum trays to carry them to the truck for exposure. All four coupons for each treatment group were placed in the same aluminum tray. To prevent cross-contamination, gloves and plastic pipettes were changed between each treatment group.

### Exposure

The aluminum trays containing the petri dishes with coupons were placed inside a truck cab (Chevrolet Z71 extended cab, General Motors, Detroit, Michigan). The trucks were housed inside a heated garage in the Iowa State University College of Veterinary Medicine Field Services Building. The truck interior was lined with a plastic sheet for each respective treatment to minimize potential contamination of the surfaces. The petri dishes were opened and placed on the truck floor in the shoe area to replicate real-world settings and allow exposure to air during treatment. The ozone machine(s) were placed inside the truck cab, along with a separate oscillating fan (Air Circulator Fan CF312, Dreo, Clifton, New Jersey) to ensure continuous air circulation and a data logger (HOBO Temperature/Relative Humidity Data Logger (MX1101 HOBO, Onset Computer Corporation, Bourne, Massachusetts) to track temperature and humidity for every treatment group every minute. For treatments involving the ozone machine, an ozone datalogger (Portable Ozone (O3) Gas Detector, GasDog, Guangxi, China) was used to monitor the concentration of ozone inside the cabin of the truck at 30-second intervals. Once the equipment was in place, the machines were plugged into power sources and turned on, the doors to the truck were closed, and the exposure time began. After the designated exposure times were completed, the petri dish lids were closed, and the samples underwent virus recovery.

### Virus recovery

A sample retrieval procedure was conducted to recover the virus from the exposed surfaces. Five milliliters of MEM solution were pipetted onto each coupon’s surface and eluted ten times over the entire surface of the coupon. Two aliquots, each with two ml of the recovered virus and MEM mixture, were transferred into two 5 ml snap cap tubes, and rapidly frozen on dry ice before being stored at -80°C until testing. To expedite the procedure, two individuals concurrently handled recovery for the four replicates of each treatment. To mitigate potential cross-contamination between treatment sets, a stringent protocol was implemented. Gloves were changed for each new treatment set to ensure no carryover of contaminants. Within each treatment set, which consisted of four replicates, a separate pipette was dedicated to each individual replicate to prevent cross-contamination. After completing all replicates within a treatment set, the plastic sheeting covering the work surface was replaced with new sheeting before proceeding to the next treatment set, thereby eliminating any potential surface contamination.

### Virus titration and immunofluorescence staining

#### Virus isolation and titration

The samples were first subjected to virus isolation in Vero cells (for PEDV) or MARC-145 cells (for PRRSV) to determine the presence of viable virus in the collected samples. Briefly, 0.2 mL of each sample was inoculated into one well of Vero cells or MARC-145 cells cultured in 24-well plates. After one-hour incubation at 37°C with 5% CO_2_, the inoculum was removed and 2 mL of the fresh post-inoculation medium was added. The culture plates were incubated at 37°C with 5% CO_2_ for 2-5 days with cytopathic effects (CPE) being checked daily. Immunofluorescence staining was conducted (see below) to verify the virus isolation outcomes. For the samples that were positive by PEDV or PRRSV virus isolation, titration was further performed. The titration of PEDV was performed using Vero cells in 96-well plates, following the method described by Chen (32). Similarly, PRRSV titration was carried out using MARC-145 cells in 96-well plates, as outlined by Neat (11). Cells in the 96-well plates were inoculated with 10-fold serially diluted PEDV or PRRSV samples (100 µL/well), with three replicate wells per dilution. Row A contained the undiluted sample, while rows B to H contained 10-fold serial dilutions prepared in post-inoculation media. The inoculated plates were incubated at 37°C with 5% CO_2_ for 1 hour. Following incubation, the inoculum was removed, and fresh post-inoculation medium specific to PEDV and PRRSV was added to the respective plates. The plates were then returned to the incubator and maintained at 37°C with 5% CO_2_ for five days. The development of cytopathic effects (CPE) was monitored daily during this period. Immunofluorescence staining was conducted to verify the virus titration results.

#### Immunofluorescence staining

The medium was removed from the plates, and the cells were washed and fixed with cold 80% acetone for 10 minutes. The fixed cell plates were air-dried, and then 50 μL/well of a 100-fold dilution of FITC-conjugated monoclonal antibody specific to the nucleocapsid protein of PEDV (SD-1F-1, Medgene Labs) or PRRSV (SDOW17-F, RTI LLC) was added to the respective plates (33,34). The plates were incubated with the antibody conjugate for 1 hour, followed by three washes with PBS, each lasting 5 minutes. The immunofluorescence staining was examined under a fluorescence microscope to identify the presence of PEDV-or PRRSV-infected cells. The end-point titer was calculated using the Reed-Muench method, with a limit of detection of 1.78 x 10 log_10_TCID_50_/ml.

### Statistical analysis

Data were analyzed separately for each virus using linear regression models in SAS ® version 9.4 (SAS Institute, Inc., Cary, North Carolina). The dependent variable was viral titer, expressed as log_10_TCID_50_/ml, while the independent variables included time, treatment type, and their interaction (time × treatment type). The treatment effects were assessed using ANOVA with a significance level of 0.05, and Tukey’s method was used to perform pairwise comparisons of least squares means (LSM) to assess differences between treatment levels.

Pearson’s correlation coefficient was applied to analyze three key relationships: 1) between ozone concentration and virus titer, with ozone concentration being the primary treatment variable of interest; 2) between ozone concentration and humidity; and 3) between ozone concentration and temperature. The rationale behind examining these correlations was to understand the potential influence of environmental factors on the efficacy of ozone treatment for virus mitigation.

## Results

All negative control replicates for both PRRSV and PEDV tested negative, confirming the absence of cross-contamination during the experimental procedures. The virus titration results for PRRSV and PEDV are summarized in Table 1, presented as the least-squares means and 95% confident intervals of log_10_TCID_50_/ml values for each machine setup.

### PRRSV

All replicates in the PRRSV treatment groups had measurable virus titers, indicating that none of the ozone treatments showed complete inactivation of the virus. The 30 g/h setup showed no significant difference in PRRSV titers—from positive controls at 30, 60, and 120 minute exposures, with least square mean values ranging from 4.56 to 4.63 log_10_TCID_50_/ml. The 38 g/h ozone treatment at 60 minutes (LSM: 3.13 log_10_TCID_50_/ml, Table 1) was the only treatment that exhibited a statistically significant reduction compared to positive controls and other ozone combinations, including the 30 and 120-minute exposures of the same treatment (LSM = 4.75 and 4.69 log_10_TCID_50_/ml, respectively). The highest 68 g/h ozone level did not differ significantly from positive controls, with LSM values ranging from 4.31 to 4.87 log_10_TCID_50_/ml.

### PEDV

The ozone treatments showed variable results (Table 1). The 30 g/h ozone treatment demonstrated significantly lower PEDV titers at 60 minutes (1.19 log_10_TCID_50_/ml) and 120 minutes (0.88 log_10_TCID_50_/ml) compared to the positive control. Additionally, the 38 g/h ozone treatment significantly reduced PEDV titers at 120 minutes (1.56 log_10_TCID_50_/ml) compared to the positive control. However, the 68 g/h ozone did not differ from positive controls at 60 and 120 minutes, and the 30-minute exposure was not significantly different from the 120-minute positive control.

### Ozone concentration and environmental factors

For all treatments, the average temperature ranged from 19.39 to 24.69°C, with the standard deviation of temperature within each treatment not exceeding 0.77°C (Table 2). The time-weighted average humidity across all treatments ranged from 29.9% to 40.4%. The lowest standard deviation of humidity within each treatment was 0.23%, while the highest was up to 3.70%.

**Table 2.**
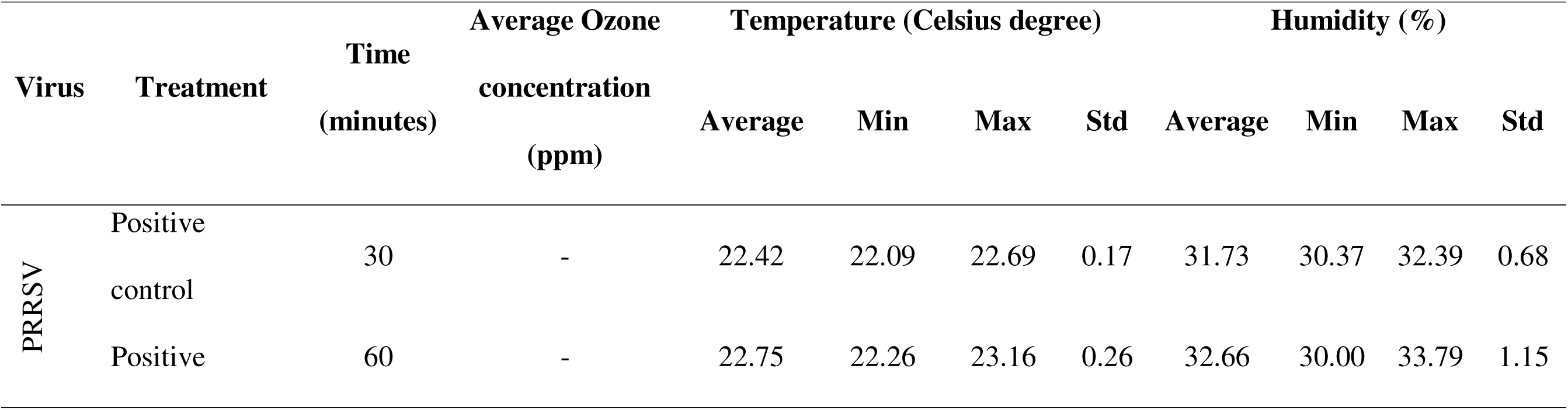
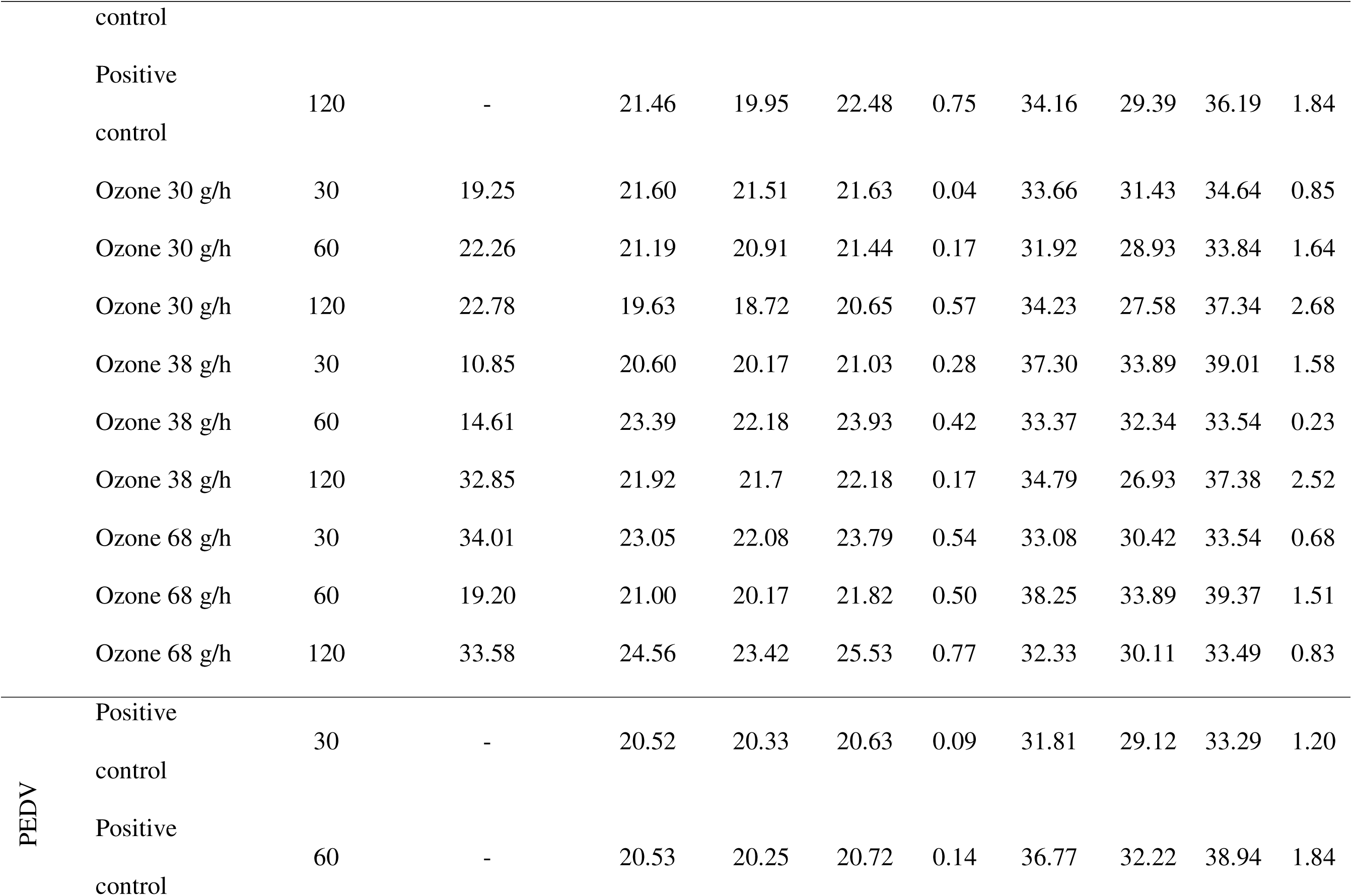
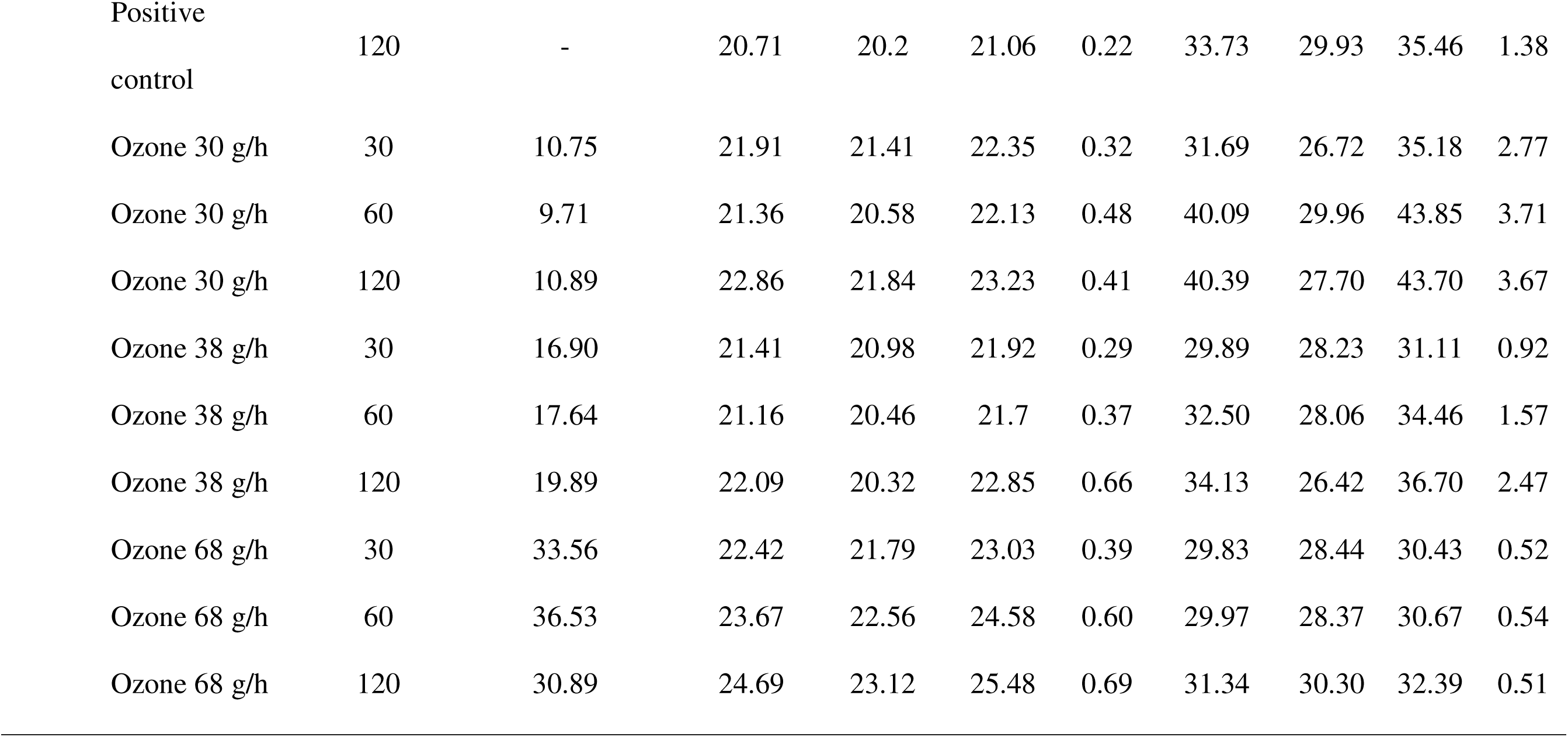
Summary of Measurements in Each Treatment: Ozone Concentration (ppm, average), Temperature (°C, Average, Max, Min, Standard Deviation), and Humidity (%, Average, Max, Min, Standard Deviation)

The results of the Pearson correlation analysis revealed a weak, statistically insignificant positive correlation between virus titer and ozone concentration for both PRRSV (r=0.21, p>0.05) and PEDV (r=0.28, p>0.05). When examining the interplay between ozone concentration and other environmental parameters, distinct patterns emerged. A moderate positive correlation was observed between ozone concentration and temperature (r=0.50, p<0.0001), suggesting that higher temperatures facilitated increased ozone generation. Conversely, a moderate negative correlation was identified between ozone concentration and humidity (r=-0.56, p<0.0001), implying that elevated humidity levels might have compromised the generation of ozone.

### Ozone concentration variability

When examining the weighted average of ozone concentration (in parts per million, ppm) during the first 30 minutes for the six treatments involving each ozone machine setup, variations in standard deviation were observed. For the same machine and duration, each machine exhibited differences in the weighted average ozone concentration. For the ozone machine with a 30 g/h capacity, the standard deviation was 4.75 ppm. The ozone machine with a 38 g/h capacity demonstrated a higher standard deviation of 0.68 ppm compared to the 30 g/h setup. When two ozone machines were combined to achieve a 68 g/h setup, the standard deviation of the weighted average ozone concentration was 5.88 ppm.

## Discussion

The results of this study demonstrate the variable efficacy of ozone gas in inactivating PRRSV and PEDV on non-porous surfaces within truck cabins at three different time levels: 30 minutes, 60 minutes, and 120 minutes. While certain ozone treatment combinations exhibited some viral reduction for both viruses, none reduced viral titers by more than two logs, which is the minimum reduction considered effective. The absence of clear trends across different exposures and concentrations suggests that the tested combinations of decontamination methods, exposure times, and ozone concentrations were insufficient for consistent and reliable inactivation of these viruses.

Notably, when compared to positive controls, more ozone gas treatments showed significant reductions in PEDV compared to PRRSV (Table 1). A study by Bolton (35) also demonstrated that even among enveloped viruses (vesicular stomatitis virus, infectious bovine rhinotracheitis virus, and influenza A virus), there can be significant differences in their responses to ozone disinfection, depending on their heat-resistant properties. At 0.64 ppm of O , all three viruses exhibited different lag periods, during which little or no effect was observed. Vesicular stomatitis virus had a short and negligible lag phase, while influenza A virus and infectious bovine rhinotracheitis virus had lag phases of 6 hours and 12 to 15 hours, respectively. At a lower O concentration of 0.16 ppm, the lag periods were even longer, with 6 hours for vesicular stomatitis virus, 8 hours for influenza A virus, and 54 hours for infectious bovine rhinotracheitis virus (35). Generally, increasing O concentration or contact time enhances disinfection efficacy (36,37), but there are thresholds beyond which further increases do not significantly improve pathogen inactivation. Insufficient O concentrations may not achieve effective inactivation (38), while excessively high concentrations offer minimal additional benefits and may result in resource inefficiency (39). Similarly, brief contact times may lead to inadequate disinfection, whereas prolonged contact times can cause a tailing effect, where the inactivation rate diminishes or stabilizes despite extended exposure (40).

However, increasing ozone concentrations can introduce other problems for users and materials. Prolonged exposure to elevated ozone levels poses potential health risks, including respiratory tract irritation and reaching toxicity thresholds, which complicates maintaining safe ozone levels for effective pathogen inactivation (41). Various organizations, such as the U.S. Environmental Protection Agency, the Occupational Safety and Health Administration, and the Food and Drug Administration, have set strict ozone exposure limits for humans at 0.08 ppm, 0.10 ppm, and 0.05 ppm, respectively, for 8 hours (42,43). Additionally, ozone application poses environmental concerns due to the potential formation of harmful byproducts from reactions with unsaturated organic compounds. These byproducts include aldehydes such as formaldehyde, acetaldehyde, and acrolein, which can be formed when ozone reacts with certain organic compounds (44,45), as well as fine and ultrafine particles that can have adverse health effects when inhaled (46). These byproducts can impact human health and air quality (41). Furthermore, ozone’s detrimental effects on surfaces, including the destruction of molecular chain networks and cross-linking points in materials like natural rubber, highlight its limitations in terms of material compatibility and long-term surface integrity (41).

The weak correlation between virus titer and ozone concentration observed in the result of this study indicates that factors other than ozone concentration affect virus inactivation. Percivalle investigated the inactivation effect of ozone gas on SARS-CoV-2 on eight different surfaces, revealing that the disinfection efficiency of gaseous ozone was not proportionate to the concentration of ozone and did not depend on the surface type (47). The study found that three concentrations of O3 gas (0.5, 1, and 2 ppm) effectively inactivated the virus within 40 minutes, and a specific ratio of humidity and hydration within droplets was crucial for the survival of SARS-CoV-2 on dry surfaces (47). These findings underscore the complexity of using ozone for viral inactivation and the need for tailored approaches based on the characteristics of each virus and the specifics of the treatment setup.

Criscuolo identified an optimal ozone concentration of 4 ppm with a 30-minute exposure to achieve over 90% inactivation of SARS-CoV-2 on various surfaces (24). In contrast, a lower, non-toxic concentration of 0.2 ppm required four times longer to achieve the same level of reduction. For truck cab decontamination, the exposure times tested in this study (30, 60, and 120 minutes) were chosen based on practical considerations. Although increasing exposure time could potentially lead to significant viral titer reductions, extending disinfection beyond 2 hours is not always feasible for field applications in truck sanitation due to impacts on vehicle utilization efficiency (48).

Environmental factors, particularly temperature and humidity, play crucial roles in influencing ozone levels and its effectiveness as a disinfectant in the air. Studies have consistently demonstrated that temperature has a positive correlation with ozone concentrations (49,50). In the context of decontamination, higher temperatures generally enhance gaseous ozone’s reactivity and decomposition rate, potentially improving its inactivation efficiency against microorganisms (51–53). Sharma and Hudson found that ozone inactivation of various viruses was more effective at 20°C than at 4°C (19). However, the relationship between temperature and ozone disinfection efficacy can vary depending on the target microorganism and other environmental factors such as humidity (18).

Relative humidity typically exhibits a negative correlation with ozone levels in the air (54–56). While higher humidity levels may decrease atmospheric ozone concentrations, humidity is generally required to achieve significant inactivation of target microorganisms during disinfection processes (18,57–60). To optimize the effect of humidity on ozone disinfection, Hudson proposed enhancing virus inactivation using a two-step process: first waiting until the measured ozone concentration reaches its maximum plateau level in the space, followed by a burst of water vapor to increase relative humidity to greater than 70% (58). A study conducted by Clavo investigated the effectiveness of ozone inactivation on pathogens found on personal protective equipment (PPE) (61). It was observed that at ozone concentrations above 2000 ppm, disinfection was achieved in less than 10 minutes, and when the concentration was increased to 10,000 ppm, disinfection was accomplished in less than 30 seconds. However, the study also revealed that at lower ozone concentrations ranging from 4-12 ppm, the effectiveness of disinfection was heavily dependent on relative humidity conditions.

The role of humidity in enhancing ozone’s activity as a disinfectant is complex. High humidity levels can enhance ozone’s oxidative properties by promoting its decomposition into highly reactive species such as hydroxyl radicals, hydrogen peroxide, and singlet oxygen, leading to the formation of more radicals than in dry air and potentially increasing virus inactivation (18,47,58,60–63). The increased efficacy observed at higher relative humidity values could also be explained by rehydration of the desiccated layer surrounding the virus, thereby enhancing ozone diffusion and virus exposure to the disinfectant (59).

Achieving acceptable disinfection rates with gaseous ozone becomes challenging at lower humidity levels, as it requires higher oxidant exposure values. Several studies have demonstrated that disinfection rates are difficult or even impossible to achieve when humidity falls below 40–50% (58–60). Notably, research conducted by Cía revealed that ozone treatments, at a concentration of 10 ppm, had no significant impact on virus viability inside emergency vehicles with low relative humidity (37%–48%) (64). While these optimal relative humidity conditions are relatively easy to achieve in a closed chamber, maintaining such high and steady relative humidity values in ventilation ducts or rooms may be challenging. High relative humidity can lead to mold growth or other damage (23), making the application of such humidity levels inside a truck cabin potentially infeasible in the long-run.

Several studies have raised doubts regarding the efficacy of gaseous ozone for virus disinfection (65–67) demonstrated that ozone application (at 140 ppm) for disinfecting HBV-contaminated hospital linen remained ineffective even with extended exposure. Lin observed limited cell mortality despite exposure to ozone concentrations ranging from 1.5 to 3 ppm (68). These findings collectively raise concerns about the reliability of ozone gas for virus disinfection, especially in contexts like emergency vehicles and hospital environments.

Limited research directly compares the effectiveness of aqueous and gaseous ozone in disinfecting enveloped viruses in the swine industry. In a study by (69) Zhang et al. (2020), they evaluated ozonized water’s efficacy against both wild-type African swine fever virus (ASFV) SY18 strain and a reporter ASFV (ASFV-ΔMGF-EGFP). They tested ozonized water at concentrations of 5, 10, and 20 mg/L for durations of 1, 3, 6, and 10 minutes at room temperature. Results showed a reduction in ASFV titers ranging from two to three log10, with a notable increase to a reduction of three log10 at 20 mg/L. They suggested ozonized water, especially at concentrations of 5 mg/L or above, may be effective for disinfection of pig farms, agricultural facilities, slaughterhouses, and meat processing plants.

### Limitations and future directions

The present study aimed to evaluate the efficacy of gaseous ozone technology for viral inactivation in truck cabins, although several limitations must be acknowledged. Despite efforts to control environmental conditions, variations in humidity and temperature across the four treatment days might have influenced the results, potentially limiting their generalizability to real-world scenarios in which conditions fluctuate both daily and seasonally. The involvement of multiple personnel in virus contamination and recovery processes might have introduced human error, and the use of commercial ozone generators from a single manufacturer limited comparisons with alternative brands or technologies. Furthermore, the experimental design, while rigorous, was limited by the use of specific rubber coupons representing truck cab flooring, which might not fully represent the diversity of surfaces found in actual vehicle interiors. The use of laboratory strains of PRRSV and PEDV at titers significantly higher than those typically encountered in field conditions, combined with the absence of organic matter in the experimental setup, potentially limits the direct applicability of these findings to real-world scenarios.

Future research should prioritize addressing these limitations to enhance the practical applicability of the findings. Evaluating the effectiveness of these technologies on a wider range of porous and non-porous materials commonly found in truck cabins would provide a more comprehensive assessment. Additionally, investigating a broader spectrum of exposure times and environmental conditions could elucidate the robustness of these decontamination methods across diverse scenarios. Exploration of potential synergies between ozone-based decontamination and other methods, such as chemical disinfectants or ultraviolet radiation, could unveil more effective approaches to viral inactivation. Assessing the impact of organic matter on decontamination efficacy, conducting comparative studies of different ozone generator brands and models, and incorporating field strains of viruses at concentrations more representative of actual contamination levels would further enhance the ecological validity of future studies. By addressing these areas, future investigations will significantly contribute to the development of improved biosecurity measures, ultimately leading to more effective and reliable decontamination protocols for vehicle interiors and similar enclosed spaces.

## Acknowledgments

This study was funded by the Swine Health Information Center (SHIC) through Project #23-033 SHIC, grant number GR-027078-00001.

