## Supplemental Table 1 for "Evaluation of gaseous ozone technology for decontaminating porcine reproductive and respiratory syndrome virus (PRRSV) and porcine epidemic diarrhea virus (PEDV) from non-porous surface in truck cabins"

**Table 1.** Least-squares means viral titers expressed as the log_10_TCID_50_/ml with 95% confident interval for each treatment group.

| **Virus** | **Time** | **Positive  controls** | **O_3_ 30 g/h  treatments** | **O_3_ 38 g/h  treatments** | **O_3_ 68 g/h  treatments** |
| --- | --- | --- | --- | --- | --- |
| PRRSV | log_10_ after 30 minutes | 4.63^bc^ [4.25, 5.00] | 4.62^bc^ [4.25, 5.00] | 4.75^abc^ [4.38, 5.12] | 4.31^c^ [3.94, 4.68] |
|  | log_10_ after 60 minutes | 4.62^bc^ [4.25, 5.00] | 4.63^bc^ [4.25, 5.00] | 3.13^d^ [2.75, 3.50] | 4.87^ab^ [4.50, 5.25] |
|  | log_10_ after 120 minutes | 5.19^a^ [4.82, 5.56] | 4.56^bc^ [4.19, 4.93] | 4.69^abc^ [4.32, 5.06] | 4.75^abc^ [4.38, 5.12] |
| PEDV | log_10_ after 30 minutes | 2.81^ab^ [2.18, 3.45] | 2.25^bcd^ [1.62, 2.88] | 2.94^ab^ [2.3, 3.57] | 1.75^cde^ [1.12, 2.38] |
|  | log_10_ after 60 minutes | 3.19^a^ [2.55, 3.82] | 1.19^e^ [0.55, 1.82] | 3.19^a^ [2.55, 3.82] | 2.75^ab^ [2.12, 3.38] |
|  | log_10_ after 120 minutes | 2.5^abc^ [1.87, 3.13] | 0.88^e^ [0.24, 1.51] | 1.56^de^ [0.93, 2.2] | 2.5^abc^ [1.87, 3.13] |

^a,b,c,d,e,f,g,h^ Different superscripts across rows and columns for each pathogen indicate statistical significance (*p*≤0.05)
