## Supplemental Table 2 for "Evaluation of gaseous ozone technology for decontaminating porcine reproductive and respiratory syndrome virus (PRRSV) and porcine epidemic diarrhea virus (PEDV) from non-porous surface in truck cabins"

**Table 2.** Summary of Measurements in Each Treatment: Ozone Concentration (ppm, average), Temperature (°C, Average, Max, Min, Standard Deviation), and Humidity (%, Average, Max, Min, Standard Deviation)

| **Virus** | **Treatment** | **Time (minutes)** | **Average Ozone concentration (ppm)** | **Temperature (Celsius degree)** | | | | **Humidity (%)** | | | |
| --- | --- | --- | --- | --- | --- | --- | --- | --- | --- | --- | --- |
|  |  |  |  | **Average** | **Min** | **Max** | **Std** | **Average** | **Min** | **Max** | **Std** |
| PRRSV | Positive control | 30 | - | 22.42 | 22.09 | 22.69 | 0.17 | 31.73 | 30.37 | 32.39 | 0.68 |
|  | Positive control | 60 | - | 22.75 | 22.26 | 23.16 | 0.26 | 32.66 | 30.00 | 33.79 | 1.15 |
|  | Positive control | 120 | - | 21.46 | 19.95 | 22.48 | 0.75 | 34.16 | 29.39 | 36.19 | 1.84 |
|  | Ozone 30 g/h | 30 | 19.25 | 21.60 | 21.51 | 21.63 | 0.04 | 33.66 | 31.43 | 34.64 | 0.85 |
|  | Ozone 30 g/h | 60 | 22.26 | 21.19 | 20.91 | 21.44 | 0.17 | 31.92 | 28.93 | 33.84 | 1.64 |
|  | Ozone 30 g/h | 120 | 22.78 | 19.63 | 18.72 | 20.65 | 0.57 | 34.23 | 27.58 | 37.34 | 2.68 |
|  | Ozone 38 g/h | 30 | 10.85 | 20.60 | 20.17 | 21.03 | 0.28 | 37.30 | 33.89 | 39.01 | 1.58 |
|  | Ozone 38 g/h | 60 | 14.61 | 23.39 | 22.18 | 23.93 | 0.42 | 33.37 | 32.34 | 33.54 | 0.23 |
|  | Ozone 38 g/h | 120 | 32.85 | 21.92 | 21.7 | 22.18 | 0.17 | 34.79 | 26.93 | 37.38 | 2.52 |
|  | Ozone 68 g/h | 30 | 34.01 | 23.05 | 22.08 | 23.79 | 0.54 | 33.08 | 30.42 | 33.54 | 0.68 |
|  | Ozone 68 g/h | 60 | 19.20 | 21.00 | 20.17 | 21.82 | 0.50 | 38.25 | 33.89 | 39.37 | 1.51 |
|  | Ozone 68 g/h | 120 | 33.58 | 24.56 | 23.42 | 25.53 | 0.77 | 32.33 | 30.11 | 33.49 | 0.83 |
| PEDV | Positive control | 30 | - | 20.52 | 20.33 | 20.63 | 0.09 | 31.81 | 29.12 | 33.29 | 1.20 |
|  | Positive control | 60 | - | 20.53 | 20.25 | 20.72 | 0.14 | 36.77 | 32.22 | 38.94 | 1.84 |
|  | Positive control | 120 | - | 20.71 | 20.2 | 21.06 | 0.22 | 33.73 | 29.93 | 35.46 | 1.38 |
|  | Ozone 30 g/h | 30 | 10.75 | 21.91 | 21.41 | 22.35 | 0.32 | 31.69 | 26.72 | 35.18 | 2.77 |
|  | Ozone 30 g/h | 60 | 9.71 | 21.36 | 20.58 | 22.13 | 0.48 | 40.09 | 29.96 | 43.85 | 3.71 |
|  | Ozone 30 g/h | 120 | 10.89 | 22.86 | 21.84 | 23.23 | 0.41 | 40.39 | 27.70 | 43.70 | 3.67 |
|  | Ozone 38 g/h | 30 | 16.90 | 21.41 | 20.98 | 21.92 | 0.29 | 29.89 | 28.23 | 31.11 | 0.92 |
|  | Ozone 38 g/h | 60 | 17.64 | 21.16 | 20.46 | 21.7 | 0.37 | 32.50 | 28.06 | 34.46 | 1.57 |
|  | Ozone 38 g/h | 120 | 19.89 | 22.09 | 20.32 | 22.85 | 0.66 | 34.13 | 26.42 | 36.70 | 2.47 |
|  | Ozone 68 g/h | 30 | 33.56 | 22.42 | 21.79 | 23.03 | 0.39 | 29.83 | 28.44 | 30.43 | 0.52 |
|  | Ozone 68 g/h | 60 | 36.53 | 23.67 | 22.56 | 24.58 | 0.60 | 29.97 | 28.37 | 30.67 | 0.54 |
|  | Ozone 68 g/h | 120 | 30.89 | 24.69 | 23.12 | 25.48 | 0.69 | 31.34 | 30.30 | 32.39 | 0.51 |
